# Dynamics of Ligand Binding Sites and Chloride Penetration in a Bitter Taste GPCR

**DOI:** 10.1101/2025.11.27.690919

**Authors:** Alon Rainish, Liel Sapir, Masha Y Niv

## Abstract

Taste perception strongly influences food choice, drug compliance, and dietary behavior. Bitter taste receptors belong to a subfamily of Family A GPCRs, while having unique features and deviations from conserved motifs. Recent CryoEM structures revealed that the human bitter taste receptor TAS2R14 contains not only the canonical extracellular binding site, but also a novel intracellular site, raising questions about their interplay. Using molecular dynamics simulations, we examined how binding of aristolochic acid in the intracellular site and cholesterol at the extracellular site, alone or together, affects receptor dynamics and pocket communication. We observed that cholesterol binding in the extracellular site expanded the intracellular pocket volume and induced conformational changes in TM6, whereas aristolochic acid binding in the intracellular site had no such effect on the extracellular site. Notably, in cholesterol-only simulations, a chloride ion entered from the intracellular side and interacted with Arginine 55 (BW position 2.50), revealing an inverted ion-binding pattern to Family A GPCRs, where aspartate 2.50 is known to bind a sodium ion to stabilize an inactive state. Our results suggest chloride-dependent and cholesterol-dependent modulation mechanisms in the dual-site dynamics of TAS2R14.

## 1. Introduction

Bitter taste receptors (TAS2Rs or T2Rs) are a subfamily of G protein coupled receptors (GPCRs) responsible for mediating the perception of bitterness (1). In humans, 25-26 functional TAS2Rs have been identified (2,3). Among them, TAS2R14 is the most broadly tuned receptor, activated by a wide range of chemically diverse ligands (4). TAS2R14 is expressed not only in the oral cavity but also in various extraoral tissues, including the heart, lungs, reproductive system and more(5). Its widespread expression suggests additional physiological roles beyond taste perception. TAS2R14 has been implicated in several disease contexts, including associations with colorectal(6) and pancreatic(7) cancers, positioning it as a potential therapeutic target.

The recent cryo-electron microscopy (cryo-EM) structures of TAS2R14 have revealed unexpected features, most notably an additional, intracellular, ligand-binding site (8–11). This ligand binding was neither predicted by computational models (12,13) nor observed in other TAS2R structures that have been solved so far, namely TAS2R46(14) and TAS2R16(15). The presence of this unusual pocket challenges traditional views of GPCR agonist binding and activation, raising questions about its functional significance and the potential interplay between intra- and extracellular environments. Recent advances in molecular dynamics have demonstrated that unbiased, multi-microsecond simulations of GPCRs can reliably sample key conformational substates and reveal ion or lipid allosteric sites (e.g., GPCRmd, which captures the dynamics of the 3D GPCR-ome)(16). To explore the behavior of this novel intracellular pocket, we applied molecular dynamics (MD) simulations to the recently solved cryo-EM structure of TAS2R14 bound to aristolochic acid (PDB:8XQO)(9), in which aristolochic acid (AA) occupies the intracellular site, while cholesterol (CLR) is bound to the extracellular site. Cholesterol has been observed at the extracellular binding site of several TAS2R14 structures(9–11), while in other structures the extracellular site was occupied by the same ligand as in intracellular site (PDB ID: 8RQL(8), 9IIX, 9IIW(11)) or with no CLR in the extracellular site at all (8XQR(9)). The presence of cholesterol raises the question regarding its potentially stabilizing or functional role at the extracellular site. Our simulations aimed to investigate the stability and behavior of the intracellular pocket and potential interplay between the intracellular and extracellular binding sites. Simulations were performed for different states, with both sites occupied, a single site occupied, and the apo state. The analysis showed that ligand at one site may affect the volume of the other site, and that the presence of cholesterol at the extracellular site is compatible with intracellular entry of a chloride ion and its binding at the TAS2R-conserved Arginine at the 2.50 Ballesteros-Weinstein (BW) position.

## 2. Materials and Methods

### 2.1. Preparation and simulation

We initially prepared four systems for the TAS2R14 MD simulations: a system of the experimental structure PDB ID: 8XQO with both ligands (herein “Both ligands”), one with only cholesterol in the extracellular site (“only CLR”), one with only aristolochic acid in the intracellular site (“only AA”) and one apo state (“apo”). These systems include the Gα subunit. An additional apo system was simulated without the Gα subunit.

All MD systems were prepared using the CHARMM-GUI Membrane Builder (17), using the histidine protonation state predicted by MolProbity (18). Missing loops were added based on the Alpha fold model (AF-Q9NYV8-F1-v6) (19). Protein preparation was performed using the Protein Preparation Wizard (Schrödinger Release 2025-1, Schrödinger, LLC, New York, NY, 2025) within the Schrödinger Suite. The complexes were oriented in the membrane using the default OPM alignment, and embedded in a palmitoyl-oleoyl-phosphatidylcholine (POPC) lipid bilayer, with a rectangular box extending at least 15 Å from the protein in each direction.

The system was solvated using CHARMM TIP3P water molecules, and 0.05 M NaCl (50 mM) was added to ensure electrostatic neutrality and to provide a moderate ionic strength. While this concentration is lower than typical extracellular Na^+^/Cl^−^ levels (∼140 mM Na^+^ and ∼100–110 mM Cl^−^), it is commonly used in MD simulations to balance computational efficiency with realistic screening of electrostatic interactions. All molecular parameters were assigned using the CHARMM36m force field for proteins and lipids (20). In ligand-bound systems, ligands were parameterized using the CHARMM General Force Field (CGenFF). The final system included the receptor, POPC lipids, water, and ions, and was exported in a format compatible with GROMACS for downstream equilibration and production simulations.

The MD simulations were performed using GROMACS 2021.5 (21). Each minimization was carried out until no further changes in potential energy were detected below a prescribed cutoff (Table S1). Six steps of equilibration, for a total of 1.875 ns were performed. In each step, the harmonic restraints applied to the protein, peptide and lipids were progressively decreased (Table S2). The temperature in each step was controlled with a v-rescale thermostat and set to 310 K, while from step 3 to 6 the pressure was set to 1 bar and controlled with semi-isotropic Berendsen barostat. Bond lengths to hydrogen atoms were constrained using the LINCS algorithm. Short range electrostatic and van der Waals interactions were cut off at 1.2 nm, while long-range electrostatic interactions were computed using the particle mesh Ewald method. Three production run replicas of 2 µs were performed for each complex with a time step of 2 fs and employing periodic boundary conditions (Table S3). The molecular dynamics were run in the NVT ensemble using the Nose-Hoover thermostat at 310K and sampling the trajectories every 50 ps.

### 2.2. Volume Calculation

Cavity volume calculations were performed using the POVME 3.0 software package (22).

### 2.3. Root Mean Square Deviation (RMSD) Analysis

RMSD analysis was used to assess the structural stability of TAS2R14 during MD simulations. Calculations were performed with Visual Molecular Dynamics (VMD) v1.9.4 using custom Tcl scripts, incorporating periodic boundary condition correction, recentering, and alignment to the reference cryo-EM structure PDB ID: 8XQO. Receptor protein backbone atoms were used for alignment and RMSD calculation. Ligand RMSD was computed based on heavy atoms after aligning each frame to the receptor backbone to assess relative movement within the binding site.

### 2.4. Root Mean Square Fluctuation (RMSF) Analysis

RMSF analysis was performed by MDAnalysis(23) (v2.0+) to quantify residue-level flexibility in TAS2R14. The simulation trajectory, corrected for periodic boundary conditions and aligned to the input structure, was analyzed for Cα atoms of the receptor. RMSF values were calculated separately for each component.

### 2.5. Water molecule count analysis

Water molecules within 3 Å of the ligand (or ligand position in ligand-free states) were counted throughout the simulation using a custom Tcl script in VMD v1.9.4.

### 2.6. PCA analysis

Conformational dynamics of TAS2R14 were analyzed by principal component analysis (PCA) on Cα atoms, using MDAnalysis for trajectory handling and coordinate extraction, and scikit-learn(24) for computing the PCA and projecting the structures onto principal component space. Trajectories were first aligned to the prepared PDB to remove overall rotation and translation, and frames were sampled every 2ns. PCA was performed on concatenated coordinates from all replicates and all four systems. Outlier frames were excluded based on a Z-score threshold of 3. Density distributions along the first two principal components were computed with Gaussian kernel density estimation, and the highest-density frames were extracted for structural inspection.

### 2.7. Distance and Chloride Occupancy Analysis

Distances between the Cα atoms of residues V230 and Y107 were measured along the trajectories using a custom tcl script in VMD, sampling every 4th frame (0.2ns) to reduce data redundancy. Chloride occupancy near the Arg 55 position was similarly quantified by identifying all chloride ions within 3 Å of Arg55 in each frame, and the fraction of frames in which an ion occupied that position was calculated. All analyses were performed across replicate trajectories for each simulation condition.

## 3. Results and discussion

To investigate the function of the intracellular binding site of TAS2R14 and its relationship with the extracellular site, we performed MD simulations using the recently solved structure of TAS2R14 (PDB: 8XQO), which includes aristolochic acid (AA) bound to the intracellular site and cholesterol (CLR) bound to the extracellular site. Four distinct receptor states were simulated, each in triplicate for 2 μs: Both ligands bound (AA and CLR), AA only (CLR removed), CLR only (AA removed) and Apo (with Gα) (Figure 1). We analyzed differences between the states focusing on both binding sites using structural and dynamic measures including volume, conformational changes, ion behavior, water distribution, movement of ligand and protein.

**Figure 1.**
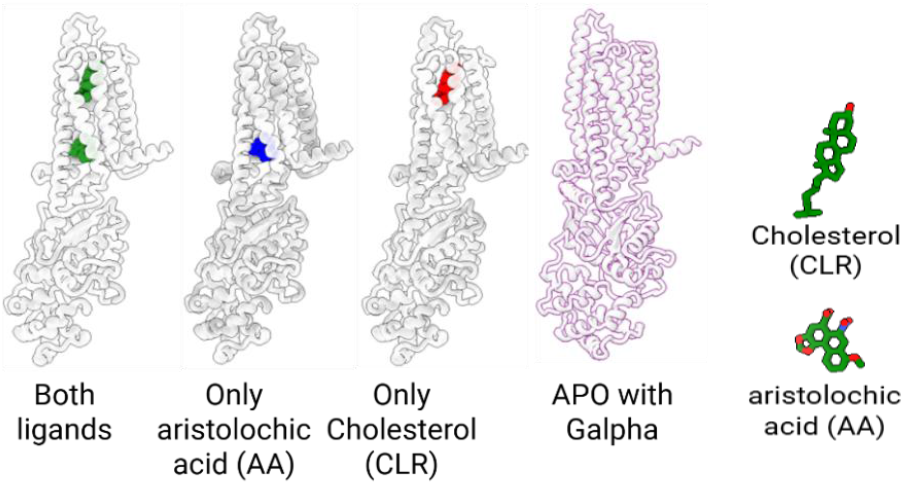
The simulated systems and the ligands bound to each of the sites. The color scheme to distinguish between the different systems is used throughout.

### 3.1. Pocket Volumes and Ligand Effects

Volume of the intracellular and extracellular binding pockets are depicted in Figure 2. States containing a bound ligand exhibited larger volumes in the extracellular site with the CLR-only state showing the greatest expansion. Cholesterol-free states displayed lower volumes with broader distributions. In the intracellular site, states with bound ligands had larger volumes, with the dual-ligand state had the largest intracellular volume, and the higher volume states are characterized by a broader distribution.

**Figure 2.**
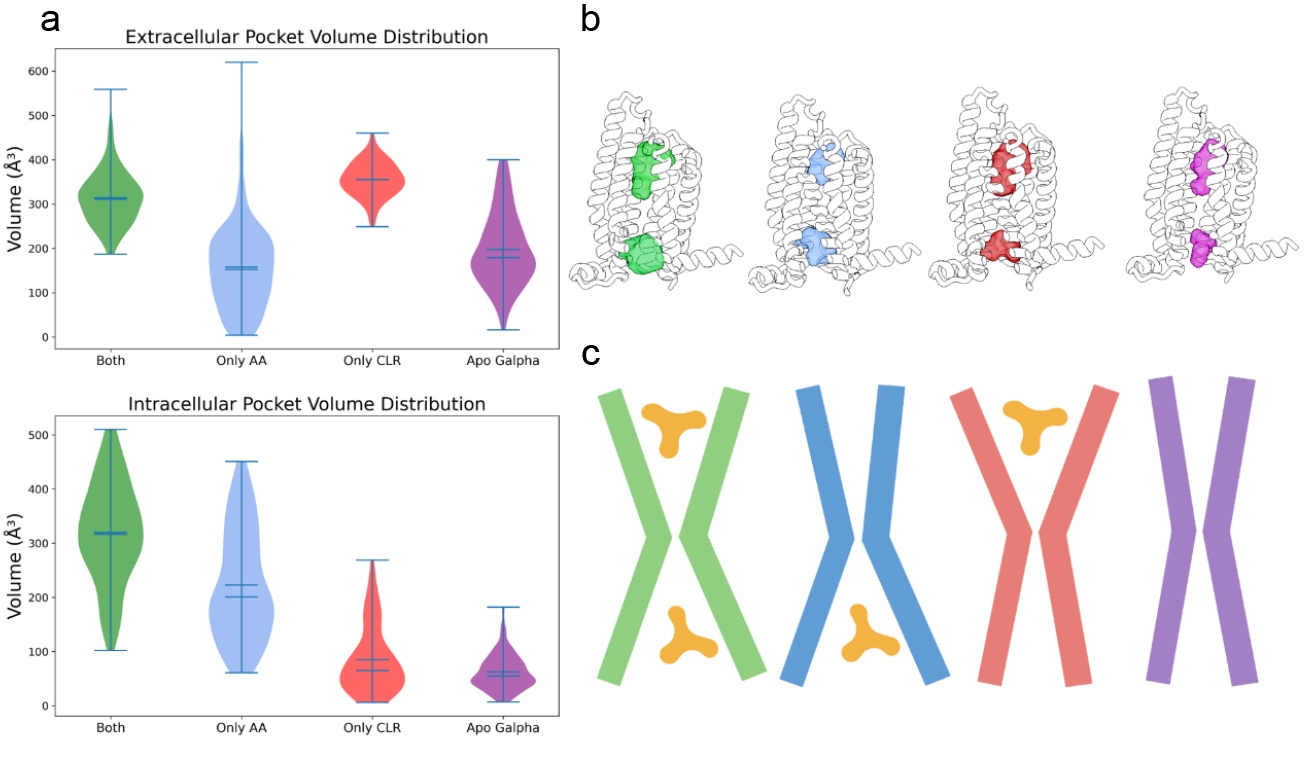
Volume analysis of the binding sites. (a) Violin plots showing the pocket volume distributions for the extracellular (top) and intracellular (bottom) binding sites across different receptor states.(b) Structural representations of the measured pocket volumes in each state, using the same color scheme: both ligands bound (green), only aristolochic acid bound at the intracellular site (blue), only cholesterol bound at the extracellular site (red), and apo state with the Gα subunit present (purple). (c) Schematic representation of the binding sites and ligand positions.

The distributions highlights differences in pocket dynamics. In the extracellular site, CLR-bound states show larger but tighter distributions, suggesting a more expanded yet restricted pocket, whereas CLR-free states are smaller but more variable, consistent with greater flexibility. In the intracellular site, AA-bound states display higher volumes with broader distributions, indicating a more open and dynamic conformation compared to the more compact AA-free states. These trends point to ligand-dependent effects on pocket flexibility. Panel (b) maps the measured volume differences onto the receptor structure, visually showing which regions of the extracellular and intracellular pockets expand in each ligand state. Panel (c) provides a simplified schematic that captures these shifts, emphasizing the overall trend of pocket opening or narrowing depending on ligand binding. We next explored the structural details underlying the volume changes.

### 3.2. Ligand-Induced Conformational Changes in TAS2R14

PCA analysis was performed on the receptor’s Cα atoms in order to evaluate allosteric changes between the states. The results indicate that the different simulated systems occupy distinct states and that the dual bound state exhibited the least diversity (Figure 3a), RMSF analysis revealed small, localized, differences, particularly in Intracellular loop 2 (ICL2), where CLR showed a stabilizing effect, extracellular loop 2 (ECL2) were CLR showed a destabilizing effect and transmembrane helix 6 (TM6) were AA showed a destabilizing effect.

**Figure 3.**
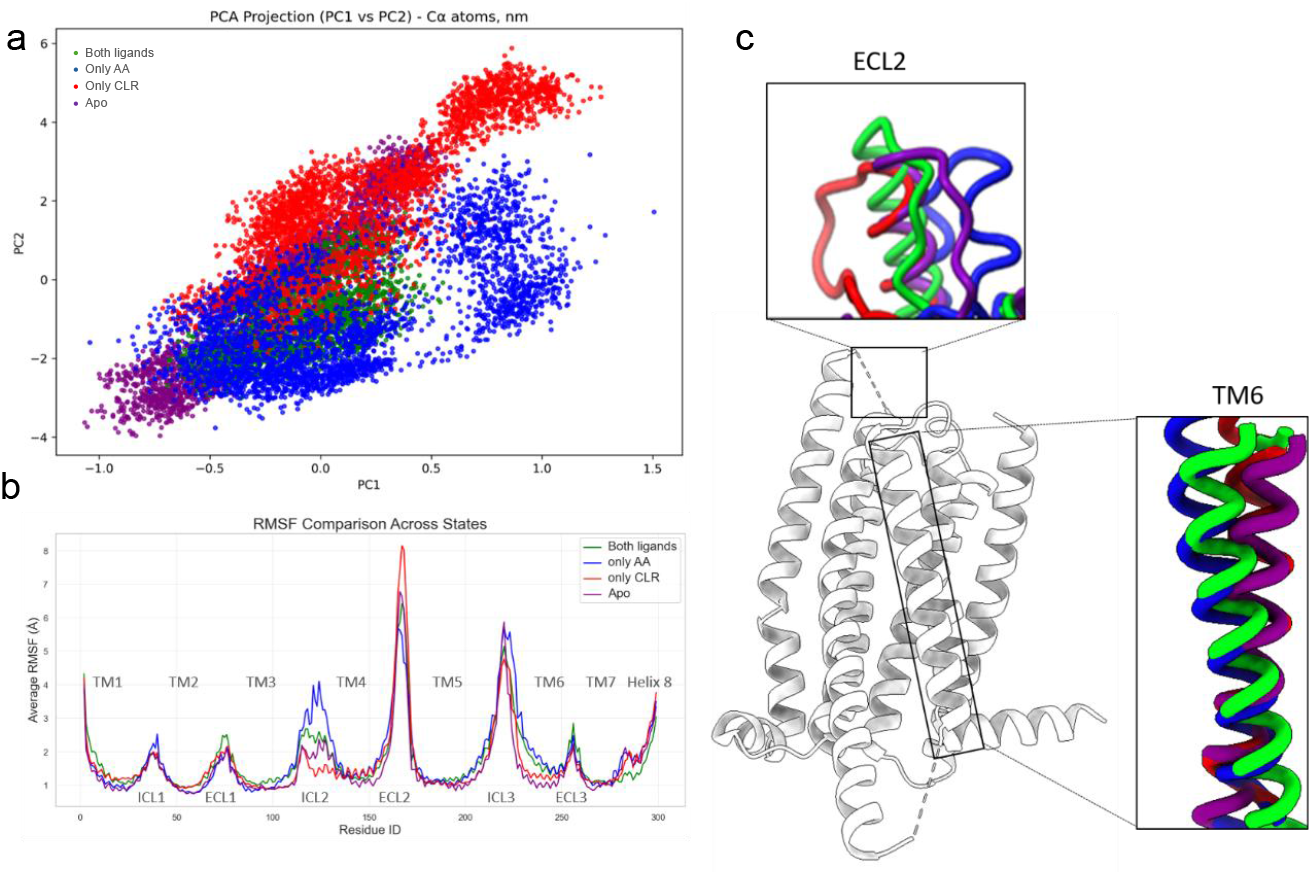
Principal component analysis (PCA) and RMSF analysis of TAS2R14 conformational dynamics. (a) PCA projections of TAS2R14 in different binding states, showing the overall conformational space and individual state distributions. (b) RMSF of the receptor for all states. Transmembrane (TM) and loop regions are annotated. (c) Structural comparison highlighting the major conformational variations between states, primarily involving movements in extracellular loop 2 (ECL2) and transmembrane helix 6 (TM6).

ECL2 is the longest and least conserved extracellular loop among class A GPCRs. It has been shown to play important roles in ligand selectivity and receptor activation, displaying distinct conformational behaviors across different receptors (25). In the studied system, ECL2, which was unresolved in the CryoEM structure, and was modelled prior to creating the different stating states for the simulations, responds differently depending on the binding profile: when CLR binds alone to the extracellular site, the loop adopts a more outwards conformation; when AA binds alone in the intracellular site, the loop moves inwards. The structure with both ligands bound shows an intermediate conformation, resembling that of the apo state but slightly more open.

Notably, TM6 movement has an important role in GPCR activation (26). In the inactive conformation of class A GPCRs, the highly conserved Arg (3.50) residue of the DRY motif typically forms a salt bridge, known as the ionic lock, with a negatively charged residue in TM6, usually Glu or Asp (6.30). This interaction holds TM6 closer to TM3, thereby stabilizing the receptor in its inactive state. Upon agonist binding, conformational rearrangements disrupt this TM3-TM6 interaction, breaking the ionic lock and allowing TM6 to swing outward in the intracellular side. This outward displacement opens the receptor’s cytoplasmic interface, enabling G protein binding and activation. PCA shows that in AA bound states, TM6 moves outwards in the bottom region and inwards at the top, unlike the states with no AA. In most GPCR classes, activation is driven by a pronounced movement of TM6. For example, in class A GPCRs, a conserved proline (6.50) induces a characteristic kink in TM6, which straightens and swings outward during activation, creating space for G-protein binding (27). In contrast, TAS2R14 exhibits a different behavior. As TAS2Rs lack this conserved proline (28), the TAS2R14 TM6 does not undergo the typical sharp kink observed in class A GPCRs. Instead, activation involves a more subtle and continuous displacement of TM6, resembling the smoother activation mechanism described for class F GPCRs(27).

### 3.3. TM6 motion and water behavior

We measured the distance between Y107 (3.50), a residue of the conserved DRY motif observed in class A GPCRs(28), and V230 (6.35). this distance is longer in the AA-bound structures, while states lacking AA display a shorter distance, consistent with a “locked” inactive-like conformation (Figure 4a). Interestingly, in the complex containing only CLR in the extracellular site, transient peaks in this distance were observed towards the end of the simulation. The late-occurring fluctuations in the TM3-TM6 opening may reflect subtle receptor activation dynamics induced by CLR binding (Figure 4), which is compatible with recent experimental data that demonstrated cholesterol to be a weak agonist of TAS2R14 (9).

**Figure 4.**
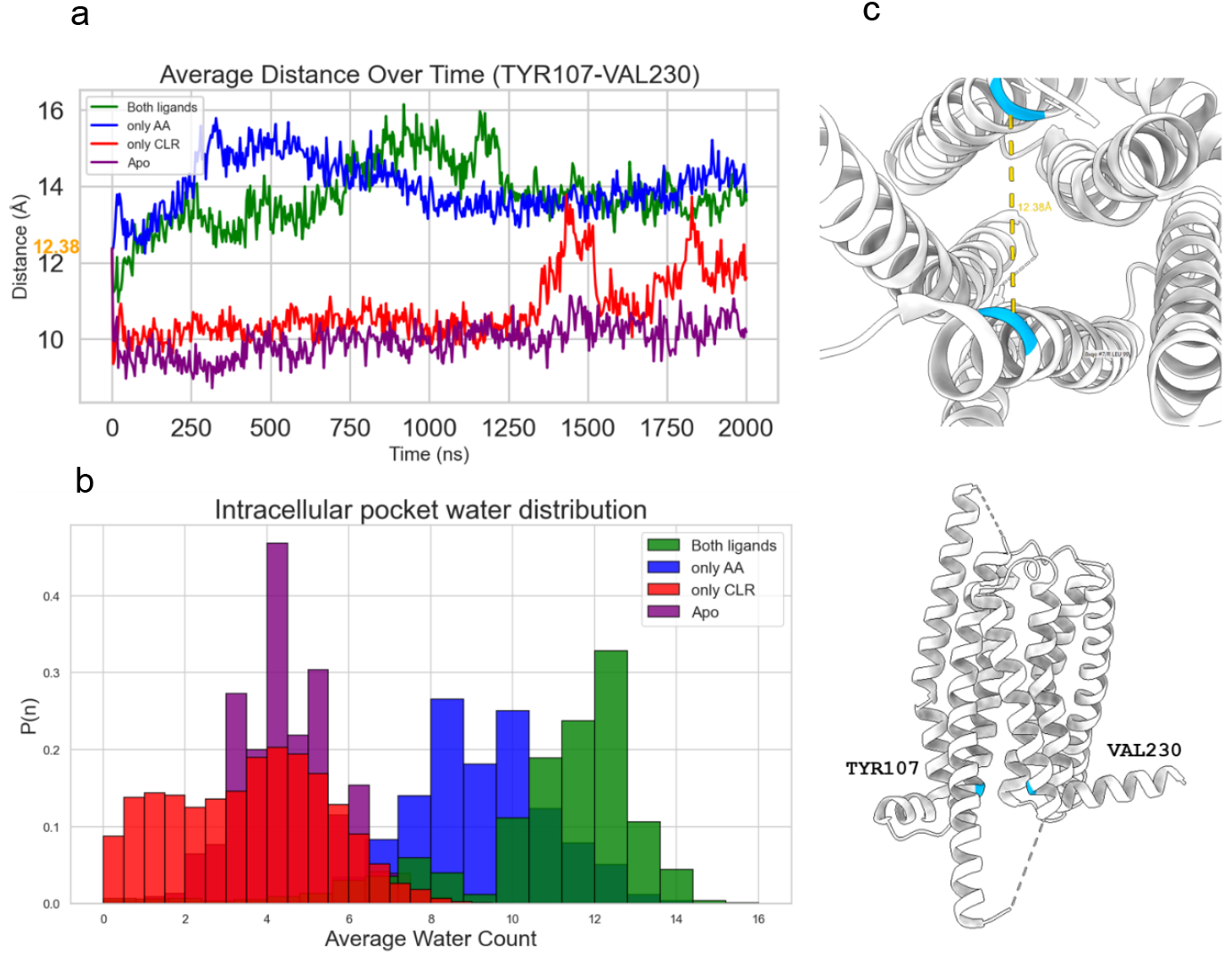
Distance between VAL230 and TYR107 in TAS2R14. (a) distance between VAL230(6.35) and TYR107(3.50) Cα’s in all different states as a function of simulation time. (b) Probability distributions of the number of water molecules within 3 Å of the ligand or equivalent empty pocket volume in the and intracellular site. (c) Location of the residues TYR107(3.50) and VAL230(6.35) and the distance in the original solved structure (8XQO), also highlighted in (a).

We next analyzed the presence of water in both the extracellular (Supplementary S1) and intracellular sites (Figure 4b). In accord with the volume and TM3-TM6 distance, complexes containing AA showed higher water occupancy in the intracellular site, with the dual-binder complex exhibiting the greatest water content. The results suggest that AA enters the site accompanied by water molecules.

### 3.4. Chlroide ion binding to TAS2R14

In class A GPCRs, a conserved sodium-binding pocket has been shown to stabilize the inactive receptor state by coordinating sodium entering from the extracellular site, with the positively charged D(2.50), polar N(7.49), and neighboring residues(29). In TAS2R14, the same positions are occupied by positively charged residues, R55 (2.50) and H276 (7.49), which are highly conserved (∼96%) across hTAS2Rs (Figure S3). We analyzed the behavior of sodium and chloride ions in our system (50 mM NaCl). While this concentration is lower than extracellular NaCl levels, it is commonly used in membrane-protein MD simulations and provides a stable ionic environment for examining ion–receptor interactions. Sodium ions did not exhibit particular interactions with the receptor in any of the simulated states, but a chloride ion entered from the intracellular side and migrated upward to interact with R55 and H276 in the absence of AA but in the presence of CLR (Figure 5a). This represents an inverted charge situation compared to Family A GPCRs. The transient occupancy of this pocket by chloride, in which a negatively charged ion arriving from the positively charged intracellular region (Figure 5b), may represent a unique modulatory mechanism of TAS2Rs.

**Figure 5.**
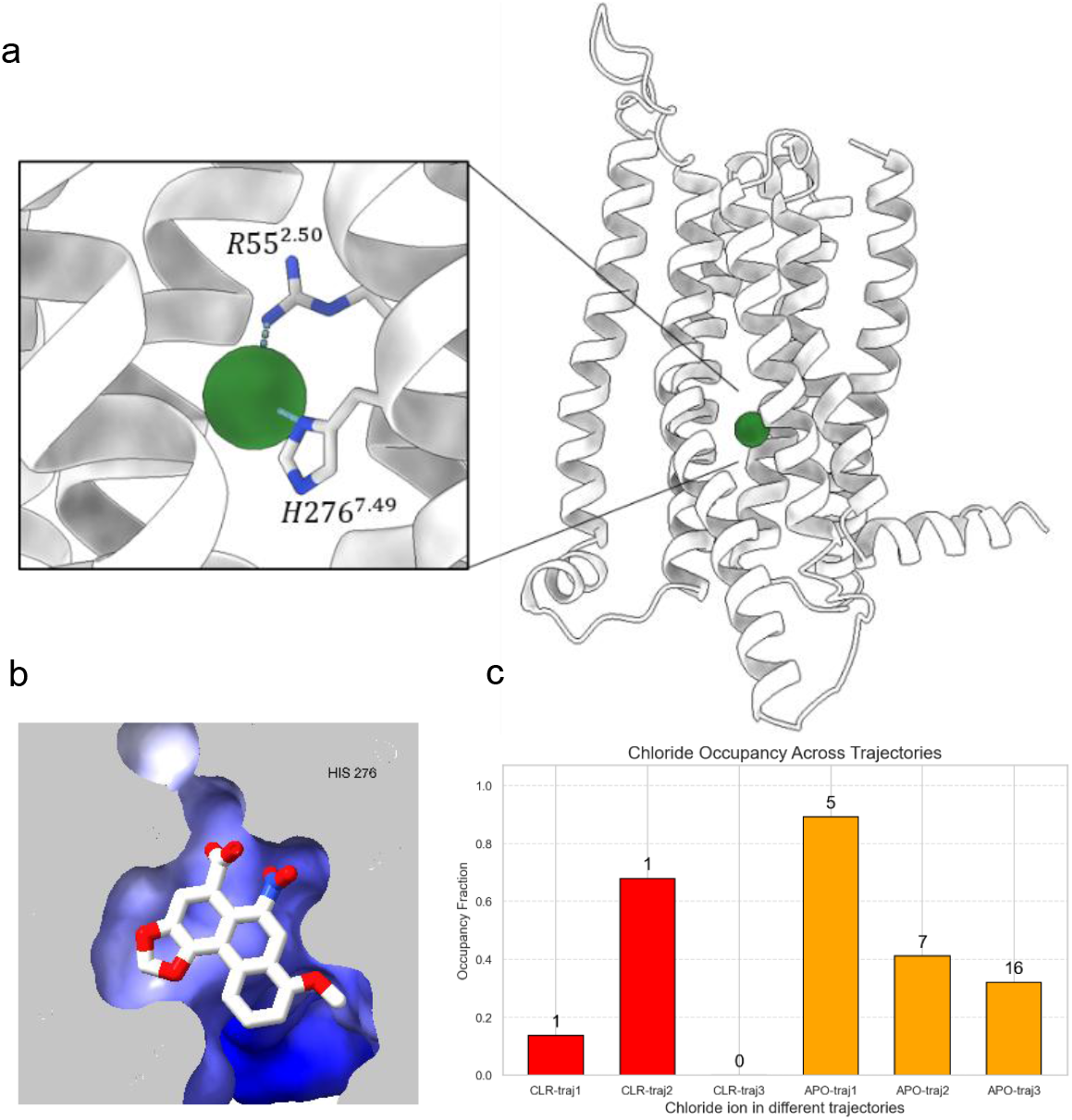
Chloride ion binding within TAS2R14. (a) Location of the chloride ion pocket and its interactions with R55 and H276. (b) AA in its intracellular pocket. The pocket is colored according to electrostatic coloring. (c) Occupancy of the potential chloride pocket throughout the simulations in which it was observed: only-CLR (red) and APO without Gα (orange). The number above each bar indicates the number of ions occupying the pocket in each simulation.

Chloride binding occurred exclusively in the CLR-only simulations and was absent in both the APO and AA-bound systems. This pattern suggests that (i) the intracellular cavity must remain sufficiently open to permit ion entry, and (ii) cholesterol facilitates such accessibility, as supported by the increasing Y107-V230 distance observed during the CLR-bound trajectories.

To further examine this behavior, we conducted additional APO simulations without Gα presence. In three independent replicas we again observed chloride entry into the same pocket. In the APO simulations without Gα, the intracellular pocket remained open, and chloride ions were repeatedly observed entering and leaving the cavity; however, they did not form stable interactions. Instead, different chloride ions transiently occupied the site, with variable residence times and overall occupancies across trajectories. In contrast, in the CLR-only state, a single chloride ion entered and remained stably bound throughout two out of three trajectories (Figure 5c).

Taken together, these findings suggest that chloride interacts with residues inside the receptor, above the newly discovered intracellular pocket. The entrance becomes accessible in specific open conformations of TAS2R14, particularly those favored by cholesterol binding, or in the absence of Gα. Notably, based on the binding patterns we observed, the role of the chloride may be more prominent when the extracellular site is occupied by CLR but the intracellular site is free. The ion’s presence in both the APO and CLR-only states indicates that chloride may only enter without intracellular ligand presence. When the entry is not blocked by a ligand, the negative ion is engaging the positively charged residues R55 and H276, the same residues that make interactions with the acidic moiety of TAS2R14 ligands (Figure 5b). A recent study experimentally checked the effects of sodium chloride on ligands that bind intracellularly (AA and flufenamic acid) and showed almost no change in activation when exposed to increasing concentration of Sodium chloride, but slightly reduced the activation of TAS2R16 by its extracellularly bound agonist salicin (30). We may speculate that the presence of chloride ion may have an affect on activation by extracellular-only ligands, while binding of intracellular acids prevents chloride from entering.

## 4. Conclusions

2μs long MD simulations revealed the flexibility of TAS2R14 binding sites, showing that both expand upon ligand binding. This property may underlie the receptor’s ability to accommodate a highly diverse set of ligands (2).

Our results suggest that the extracellular ligand influences the intracellular site, as shown by the conformational coupling between binding pockets and the formation or disruption of the canonical “lock” between TM3 and TM6. Since cholesterol has been suggested to act as a weak agonist (8), it is reasonable to propose that its binding induces partial opening of the intracellular region without enabling full receptor activation. This is further evidenced by our PCA and RMSF analyses, which reveal that conformational differences between binding states are most pronounced in ECL2 and TM6. Notably, TM6 adopts opening patterns distinct from those seen in most Class A GPCRs, particularly in the presence of the intracellular agonist aristolochic acid. The AA-bound simulations consistently show increased TYR107-VAL230 distances, indicating that ligand occupation at the intracellular site locally promotes opening of the cytoplasmic face of the receptor (28).

Further insight came from analysis of chloride ions behavior. Chloride binding occurred exclusively in the CLR-only simulations and was absent in both the APO+Gα and AA-bound systems. Combined with the observed increase in the TYR107-VAL230 distance and increased pocket volume, this suggests that cholesterol facilitates opening of the intracellular cavity, allowing ion entry. This is interesting in view of the observations reported for Class A GPCRs, where sodium helps stabilize the inactive state(29).

In the human body, chloride has a higher concentration in the extra cellular matrix than in the cytosol, which makes the observed entrance of the ion from the intracellular site even more interesting. Moreover, the chloride concentration is different in different organelles inside the cell, and the control of concentrations is mostly done by ion channels (31). Our findings indicate ligand-dependent communication between TAS2R14 binding sites, and a charge-inverted (compared to Family A) ion binding site at position 2.50 . The observed dynamics opens new questions on the role of negative ions in the intracellular environment and their potential influence on receptor function.

## Authors contributions

AR: simulations, results analysis, visualization, writing of the original draft, LS: supervision, methodology and review & editing of the draft. MYN: Conceptualization, supervision, resources, review & editing of the draft

## Acknowledgements

MYN is supported by the ISF grants 1129/19 and 1096/25. AR is a recipient of the Kennedy Leigh-Lavin PhD Scholarship Program in Sustainable Food Systems. AR is part of #CA22161 and #CA21160. MYN is part of #CA22161, #CA20128 and #CA21160. Support from FLavoursome COST action #CA22161, including STSM scientific visit and training school are gratefully acknowledged. The authors thank Alessandro Nicoli, Antonella Di Pizio, the Di Pizio lab members, Adrián García Recio and the rest of the GPCRMD team, and Fabrizio Fierro and Michael Naim for fruitful discussions.

## Supplementary information for

### simulation steps

**Table S1.** minimization steps.

| step | nsteps | Integrator | emtol<br>(kJ·mol<sup>-1</sup>) | Pos. restraints<br>BB / SC / Lipid<br>(kJ·mol<sup>-1</sup>·nm<sup>-2</sup>) | DIHRES | DIHRES_FC<br>(kJ·mol<sup>-1</sup>·nm<sup>-2</sup>) | Constraint |
| --- | --- | --- | --- | --- | --- | --- | --- |
| 1 | 5000 | steep | 1000.0 | 4000 / 2000 / 1000 | yes | 1000.0 | h-bonds (LINCS) |
| 2 | 50000 | steep | 0.001 | 2000 / 1000 / 500 | yes | 500.0 | h-bonds (LINCS) |
| 3 | 50000 | steep | 0.001 | 250 / 500 / 1000 | yes | 250.0 | h-bonds (LINCS) |
| 4 | 50000 | steep | 0.001 | 0 / 250 / 500 | yes | 0.0 | h-bonds (LINCS) |
| 5 | 50000 | steep | 0.001 | 0 / 0 / 0 | yes | 0.0 | h-bonds (LINCS) |

-LINCS was used for all hydrogen bond constraints throughout the simulation.
-Pos. restraints: positional restraints applied to backbone (BB), sidechains (SC), and lipids.
-DIHRES: dihedral restraints applied (yes/no).
-DIHRES FC: force constant for dihedral restraints.
-All abbreviations are defined in the column headers; energy units are kJ·mol<sup>-1</sup> and distances in nm<sup>-2</sup>.

**Table S2.** equilibration steps.

| step | BB restr.<br>(KJ/mol*nm2) | SC restr.<br>(KJ/mol*nm2) | Lipid restr.<br>(kJ/mol*nm2) | Thermostat | Barostat | Timestep<br>(fs) | Duration<br>(ns) |
| --- | --- | --- | --- | --- | --- | --- | --- |
| 1 | 4000 | 2000 | 1000 | v-rescale | / | 1 | 0.125 |
| 2 | 2000 | 1000 | 400 | v-rescale | / | 1 | 0.125 |
| 3 | 1000 | 500 | 400 | v-rescale | C-rescale | 1 | 0.125 |
| 4 | 500 | 200 | 200 | v-rescale | C-rescale | 2 | 0.5 |
| 5 | 200 | 50 | 40 | v-rescale | C-rescale | 2 | 0.5 |
| 6 | 50 | 0 | 0 | v-rescale | C-rescale | 2 | 0.5 |

**Table S3.** trajectory description.

| Trajectory | Galpha subunit | Ligand | Length(μs) |
| --- | --- | --- | --- |
| TAS2R14 (PDB: 8XQO) | V | cholesterol and Aristolochic acid | 2 |
| TAS2R14 (PDB: 8XQO) | V | Cholesterol | 2 |
| TAS2R14 (PDB: 8XQO) | V | Aristolochic acid | 2 |
| TAS2R14 (PDB: 8XQO) | V | - | 2 |
| TAS2R14 (PDB: 8XQO) | X | - | 2 |

BB=Backbone, SC=sidechain, restr=restrain

### Povme coordinates

For the extracellular site: A spherical inclusion region with a radius of 4 Å was defined around the center of the pocket to enclose the binding site. The grid spacing was set to 1.0 Å, and a distance cutoff of 1.09 Å was used to determine inclusion of grid points within the pocket. For the intracellular site: A spherical inclusion region with a radius of 6 Å was defined around the center of the pocket to enclose the binding site. The grid spacing was set to 1.0 Å, and a distance cutoff of 1.09 Å was used to determine inclusion of grid points within the pocket. Volumes were computed for every 200 frames of the simulation trajectory, and the resulting data were averaged across frames to assess pocket dynamics over time.

### Water Evaluation

We were able to count a large number of molecules in the extracellular site in the states with no cholesterol present, while in the intracellular site, water molecules are in higher numbers in the states with AA in the pocket (Figure S1). These opposing trends suggest that different hydration patterns govern each site’s dynamics and ligand interactions.

**Figure S1.**
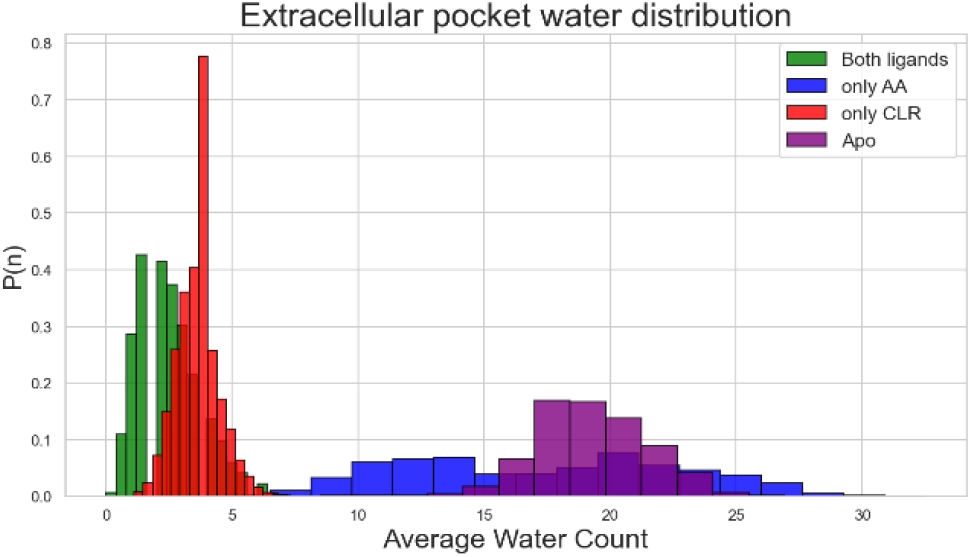
Water molecule count in the extracellular site. Number of water molecules within 3 Å of the ligand or equivalent empty pocket volume in the extracellular site. Colors correspond to the general color scheme.

This pattern suggests that water molecules within the binding sites contribute to the binding profile of each site.

### 3.3. RMSD

RMSD was calculated for each ligand using the receptor as the reference structure. In the intracellular site, AA exhibited lower RMSD when bound alone, indicating greater stability in the absence of CLR. When CLR was present in the extracellular site, AA’s RMSD increased, suggesting that ligand occupancy in the opposite site enhances AA’s mobility rather than stabilizing it (fig S2.a) towards the end in one of the only-AA trajectories, AA was able to leave the site which explains the hump in RMSD. A similar trend was observed in the extracellular site: CLR was more stable when bound alone, while its RMSD increased when AA was simultaneously bound intracellularly (fig S2.b).

Cα RMSD analysis indicated comparable overall stability across receptor states, with the dual-ligand state showing slightly higher stability over time (Fig. S2c). Structures were aligned to their reference conformation, so receptor RMSD reflects internal conformational changes. Ligand RMSD, calculated with the same alignment, primarily captures translation and rotation within the binding site rather than conformational changes of the ligands themselves. These results indicate that ligand-binding status affects ligand positioning within the pockets but has minimal impact on overall receptor stability.

**Figure S2.**
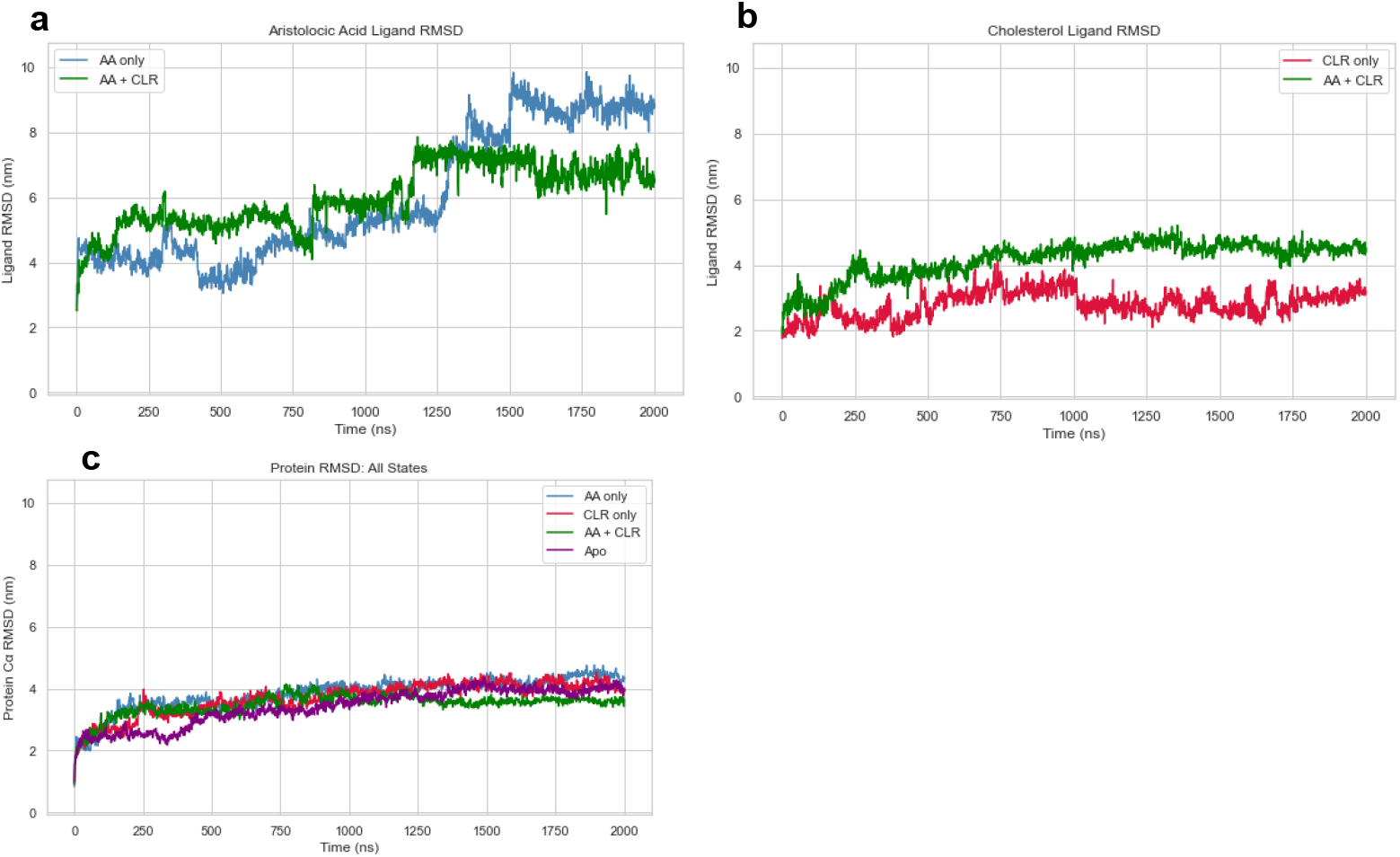
RMSD analysis of TAS2R14 receptor and ligands. (a) RMSD of AA when bound alone (blue) or together with CLR in the extracellular site (green). (b) RMSD of CLR when bound alone (red) or together with AA in the intracellular site (green). (c) Cα RMSD of the receptor in all simulated states: AA only (blue), CLR only (red), both ligands bound (green), and apo with Gα subunit (purple).

**Figure S3.**
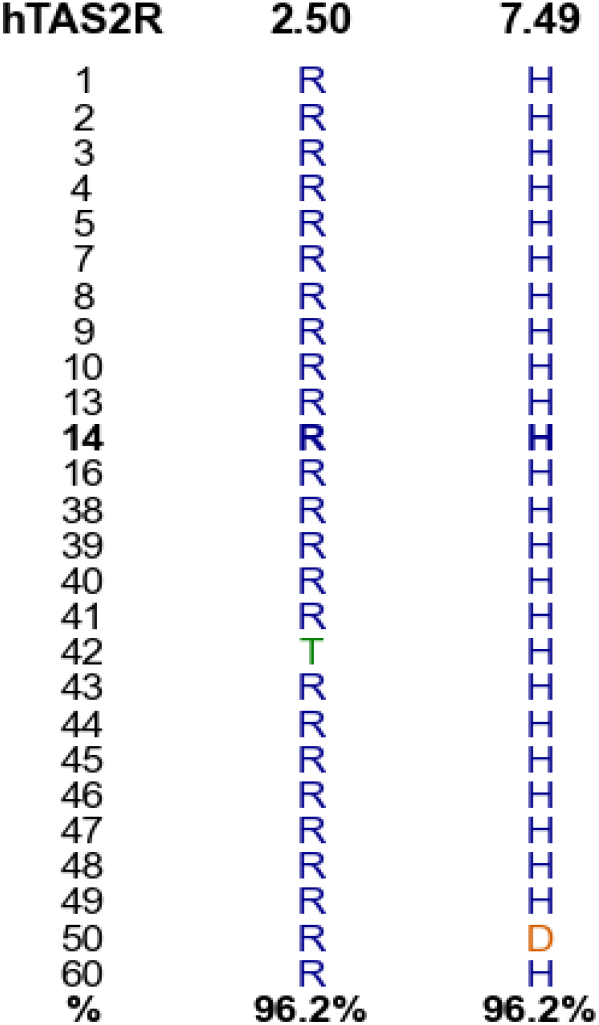
Conservation of positions 2.50 and 7.49 across human TAS2Rs. Multiple sequence alignment of human TAS2Rs showing residues at positions 2.50 and 7.49 (Ballesteros-Weinstein numbering). Each row represents a receptor, and the bottom row indicates percentage conservation.

